## Supplemental File 5 for "Analysis of promoter-associated chromatin interactions reveals biologically relevant candidate target genes at endometrial cancer risk loci"

**Endometrial Cancer Association Consortium**

Tracy A. O'Mara^1^, Deborah J. Thompson^2^, Amanda B. Spurdle^1^, Frederic Amant^3^, Daniela Annibali^3^, Katie Ashton^4-6^, John Attia^4, 7^, Paul L. Auer^8, 9^, Matthias W. Beckmann^10^, Amanda Black^11^, Louise Brinton^11^, Daniel D. Buchanan^12-15^, Stephen J. Chanock^16^, Chu Chen^17^, Maxine M. Chen^18^, Timothy H.T. Cheng^19^, Linda S. Cook^20, 21^, Marta Crous-Bous^18, 22^, Kamila Czene^23^, Immaculata De Vivo^18, 22^, Jeroen Depreeuw^3, 24, 25^, Jennifer Anne Doherty^26^, Thilo Dörk^27^, Sean C. Dowdy^28^, Alison M. Dunning^29^, Matthias Dürst^30^, Douglas F. Easton^2, 29^, Arif B. Ekici^31^, Peter A. Fasching^10, 32^, Brooke L. Fridley^33^, Christine M. Friedenreich^21^, Montserrat García-Closas^16, 34^, Mia M. Gaudet^35^, Graham G. Giles^13, 36, 56^, Dylan M. Glubb^1^, Ellen L. Goode^37^, Maggie Gorman^19^, Christopher A. Haiman^38^, Per Hall^23, 39^, Susan E. Hankinson^22, 40^, Patricia Hartge^11^, Catherine S. Healey^29^, Alexander Hein^10^, Peter Hillemanns^27^, Shirley Hodgson^41^, Erling Hoivik^42, 43^, Elizabeth G. Holliday^4, 7^, David J. Hunter^18, 44, 45^, Angela Jones^19^, Peter Kraft^18, 44^, Camilla Krakstad^42, 43^, Diether Lambrechts^25, 46^, Loic Le Marchand^47^, Xiaolin Liang^48^, Annika Lindblom^49, 50^, Jolanta Lissowska^51^, Jirong Long^52^, Lingeng Lu^53^, Anthony M. Magliocco^54^, Lynn Martin^55^, Mark McEvoy^7^, Roger L. Milne^13, 36, 56^, Miriam Mints^57^, Rami Nassir^58^, Irene Orlow^48^, Geoffrey Otton^59^, Claire Palles^19^, Paul D.P. Pharoah^2, 29^, Loreall Pooler^38^, Jennifer Prescott^22^, Tony Proietto^59^, Timothy R. Rebbeck^60, 61^, Stefan P. Renner^62^, Harvey A. Risch^53^, Matthias Rübner^62^, Ingo Runnebaum^30^, Carlotta Sacerdote^63, 64^, Gloria E. Sarto^65^, Fredrick Schumacher^66^, Rodney J. Scott^4, 6, 67^, V. Wendy Setiawan^38^, Mitul Shah^29^, Xin Sheng^38^, Xiao-Ou Shu^52^, Melissa C. Southey^12, 56^, Emma Tham^49, 68^, Ian Tomlinson^19, 55^, Jone Trovik^42, 43^, Constance Turman^18^, David Van Den Berg^38^, Adriaan Vanderstichele^69^, Zhaoming Wang^11^, Penelope M. Webb^70^, Nicolas Wentzensen^11^, Stacey J. Winham^71^, Lucy Xia^38^, Yong-Bing Xiang^72^, Hannah P. Yang^11^, Herbert Yu^47^, Wei Zheng^52^

^1^ Department of Genetics and Computational Biology, QIMR Berghofer Medical Research Institute, Brisbane, Queensland, Australia.

^2^ Centre for Cancer Genetic Epidemiology, Department of Public Health and Primary Care, University of Cambridge, Cambridge, UK.

^3^ Department of Obstetrics and Gynecology, Division of Gynecologic Oncology, University Hospitals KU Leuven, University of Leuven, Leuven, Belgium.

^4^ Hunter Medical Research Institute, John Hunter Hospital, Newcastle, New South Wales, Australia.

^5^ Centre for Information Based Medicine, University of Newcastle, Callaghan, New South Wales, Australia.

^6^ Discipline of Medical Genetics, School of Biomedical Sciences and Pharmacy, Faculty of Health, University of Newcastle, Callaghan, New South Wales, Australia.

^7^ Centre for Clinical Epidemiology and Biostatistics, School of Medicine and Public Health, University of Newcastle, Callaghan, New South Wales, Australia.

^8^ Cancer Prevention Program, Fred Hutchinson Cancer Research Center, Seattle, WA, USA.

^9^ Zilber School of Public Health, University of Wisconsin-Milwaukee, Milwaukee, WI, USA.

^10^ Department of Gynecology and Obstetrics, Comprehensive Cancer Center ER-EMN, University Hospital Erlangen, Friedrich-Alexander-University Erlangen-Nuremberg, Erlangen, Germany.

^11^ Division of Cancer Epidemiology and Genetics, National Cancer Institute, Bethesda, MD, USA.

^12^ Department of Clinical Pathology, The University of Melbourne, Melbourne, Victoria, Australia.

^13^ Centre for Epidemiology and Biostatistics, Melbourne School of Population and Global Health, The University of Melbourne, Melbourne, Victoria, Australia.

^14^ Genomic Medicine and Family Cancer Clinic, Royal Melbourne Hospital, Parkville, Victoria, Australia.

^15^ University of Melbourne Centre for Cancer Research, Victorian Comprehensive Cancer Centre, Parkville, Victoria, Australia.

^16^ Division of Cancer Epidemiology and Genetics, National Cancer Institute, National Institutes of Health, Department of Health and Human Services, Bethesda, MD, USA.

^17^ Epidemiology Program, Fred Hutchinson Cancer Research Center, Seattle, WA, USA.

^18^ Department of Epidemiology, Harvard T.H. Chan School of Public Health, Boston, MA, USA.

^19^ Wellcome Trust Centre for Human Genetics and Oxford NIHR Biomedical Research Centre, University of Oxford, Oxford, UK.

^20^ University of New Mexico Health Sciences Center, University of New Mexico, Albuquerque, NM, USA.

^21^ Department of Cancer Epidemiology and Prevention Research, Alberta Health Services, Calgary, AB, Canada.

^22^ Channing Division of Network Medicine, Department of Medicine, Brigham and Women's Hospital and Harvard Medical School, Boston, MA, USA.

^23^ Department of Medical Epidemiology and Biostatistics, Karolinska Institutet, Stockholm, Sweden.

^24^ Vesalius Research Center, VIB, Leuven, Belgium.

^25^ Laboratory for Translational Genetics, Department of Human Genetics, University of Leuven, Leuven, Belgium.

^26^ Huntsman Cancer Institute, Department of Population Health Sciences, University of Utah, Salt Lake City, UT, USA.

^27^ Gynaecology Research Unit, Hannover Medical School, Hannover, Germany.

^28^ Department of Obstetrics and Gynecology, Division of Gynecologic Oncology, Mayo Clinic, Rochester, MN, USA.

^29^ Centre for Cancer Genetic Epidemiology, Department of Oncology, University of Cambridge, Cambridge, UK.

^30^ Department of Gynaecology, Jena University Hospital - Friedrich Schiller University, Jena, Germany.

^31^ Institute of Human Genetics, University Hospital Erlangen, Friedrich-Alexander University Erlangen-Nuremberg, Comprehensive Cancer Center Erlangen-EMN, Erlangen, Germany.

^32^ David Geffen School of Medicine, Department of Medicine Division of Hematology and Oncology, University of California at Los Angeles, Los Angeles, CA, USA.

^33^ Department of Biostatistics, Kansas University Medical Center, Kansas City, KS, USA.

^34^ Division of Genetics and Epidemiology, Institute of Cancer Research, London, UK.

^35^ Behavioral and Epidemiology Research Group, American Cancer Society, Atlanta, GA, USA.

^36^ Cancer Epidemiology Division, Cancer Council Victoria, Melbourne, Victoria, Australia.

^37^ Department of Health Science Research, Division of Epidemiology, Mayo Clinic, Rochester, MN, USA.

^38^ Department of Preventive Medicine, Keck School of Medicine, University of Southern California, Los Angeles, CA, USA.

^39^ Department of Oncology, Södersjukhuset, Stockholm, Sweden.

^40^ Department of Biostatistics & Epidemiology, University of Massachusetts, Amherst, Amherst, MA, USA.

^41^ Department of Clinical Genetics, St George's, University of London, London, UK.

^42^ Centre for Cancerbiomarkers, Department of Clinical Science, University of Bergen, Bergen, Norway.

^43^ Department of Gynecology and Obstetrics, Haukeland University Hospital, Bergen, Norway.

^44^ Program in Genetic Epidemiology and Statistical Genetics, Harvard T.H. Chan School of Public Health, Boston, MA, USA.

^45^ Nuffield Department of Population Health, University of Oxford, Oxford, UK.

^46^ VIB Center for Cancer Biology, VIB, Leuven, Belgium.

^47^ Epidemiology Program, University of Hawaii Cancer Center, Honolulu, HI, USA.

^48^ Department of Epidemiology and Biostatistics, Memorial Sloan-Kettering Cancer Center, New York, NY, USA.

^49^ Department of Molecular Medicine and Surgery, Karolinska Institutet, Stockholm, Sweden.

^50^ Department of Clinical Genetics, Karolinska University Hospital, Stockholm, Sweden.

^51^ Department of Cancer Epidemiology and Prevention, M. Sklodowska-Curie Cancer Center, Oncology Institute, Warsaw, Poland.

^52^ Division of Epidemiology, Department of Medicine, Vanderbilt Epidemiology Center, Vanderbilt-Ingram Cancer Center, Vanderbilt University School of Medicine, Nashville, TN, USA.

^53^ Chronic Disease Epidemiology, Yale School of Public Health, New Haven, CT, USA.

^54^ Department of Anatomic Pathology, Moffitt Cancer Center & Research Institute, Tampa, FL, USA.

^55^ Institute of Cancer and Genomic Sciences, University of Birmingham, Birmingham, UK.

^56^ Precision Medicine, School of Clinical Sciences at Monash Health, Monash University, Clayton, Victoria, Australia.

^57^ Department of Women's and Children's Health, Karolinska Institutet, Stockholm, Sweden.

^58^ Department of Biochemistry and Molecular Medicine, University of California Davis, Davis, CA, USA.

^59^ School of Medicine and Public Health, University of Newcastle, Callaghan, New South Wales, Australia.

^60^ Harvard T.H. Chan School of Public Health, Boston, MA, USA.

^61^ Dana-Farber Cancer Institute, Boston, MA, USA.

^62^ Department of Gynaecology and Obstetrics, University Hospital Erlangen, Friedrich-Alexander University Erlangen-Nuremberg, Comprehensive Cancer Center Erlangen-EMN, Erlangen, Germany.

^63^ Center for Cancer Prevention (CPO-Peimonte), Turin, Italy.

^64^ Human Genetics Foundation (HuGeF), Turino, Italy.

^65^ Department of Obstetrics and Gynecology, School of Medicine and Public Health, University of Wisconsin, Madison, WI, USA.

^66^ Department of Population and Quantitative Health Sciences, Case Western Reserve University, Cleveland, OH, USA.

^67^ Division of Molecular Medicine, Pathology North, John Hunter Hospital, Newcastle, New South Wales, Australia.

^68^ Clinical Genetics, Karolinska Institutet, Stockholm, Sweden.

^69^ Division of Gynecologic Oncology, Department of Obstetrics and Gynaecology and Leuven Cancer Institute, University Hospitals Leuven, Leuven, Belgium.

^70^ Population Health Department, QIMR Berghofer Medical Research Institute, Brisbane, Queensland, Australia.

^71^ Department of Health Science Research, Division of Biomedical Statistics and Informatics, Mayo Clinic, Rochester, MN, USA.

^72^ State Key Laboratory of Oncogene and Related Genes & Department of Epidemiology, Shanghai Cancer Institute, Renji Hospital, Shanghai Jiaotong University School of Medicine, Shanghai, China.

**ECAC Acknowledgements**

The authors thank the many individuals who participated in this study and the numerous institutions and their staff who supported recruitment.

The iCOGS and OncoArray endometrial cancer analysis were supported by NHMRC project grants [ID#1031333 & ID#1109286] Funding for the iCOGS infrastructure came from: the European Community's Seventh Framework Programme under grant agreement no 223175 [HEALTH-F2-2009-223175] [COGS], Cancer Research UK [C1287/A10118, C1287/A 10710, C12292/A11174, C1281/A12014, C5047/A8384, C5047/A15007, C5047/A10692, C8197/A16565], the National Institutes of Health [CA128978] and Post-Cancer GWAS initiative [1U19 CA148537, 1U19 CA148065 and 1U19 CA148112 - the GAME-ON initiative], the Department of Defence [W81XWH-10-1-0341], the Canadian Institutes of Health Research [CIHR] for the CIHR Team in Familial Risks of Breast Cancer, Komen Foundation for the Cure, the Breast Cancer Research Foundation, and the Ovarian Cancer Research Fund. OncoArray genotyping of ECAC cases was performed with the generous assistance of the Ovarian Cancer Association Consortium (OCAC). We particularly thank the efforts of Cathy Phelan. The OCAC OncoArray genotyping project was funded through grants from the US National Institutes of Health (CA1X01HG007491-01, U19-CA148112, R01-CA149429 and R01-CA058598); Canadian Institutes of Health Research (MOP-86727); and the Ovarian Cancer Research Fund. CIDR genotyping for the Oncoarray was conducted under contract 268201200008I. OncoArray genotyping of the BCAC controls was funded by Genome Canada Grant GPH-129344, NIH Grant U19 CA148065, and Cancer UK Grant C1287/A16563.

Stage 1 and stage 2 case genotyping was supported by the NHMRC [ID#552402, ID#1031333]. Control data were generated by the Wellcome Trust Case Control Consortium (WTCCC), and a full list of the investigators who contributed to the generation of the data is available from the WTCCC website. We acknowledge use of DNA from the British 1958 Birth Cohort collection, funded by the Medical Research Council grant G0000934 and the Wellcome Trust grant 068545/Z/02 - funding for this project was provided by the Wellcome Trust under award 085475. NSECG was supported by the EU FP7 CHIBCHA grant, Wellcome Trust Centre for Human Genetics Core Grant 090532/Z/09Z, and CORGI was funded by Cancer Research UK. We thank Nick Martin, Dale Nyholt and Anjali Henders for access to GWAS data from QIMR Controls. Recruitment of the QIMR controls was supported by the NHMRC. The University of Newcastle, the Gladys M Brawn Senior Research Fellowship scheme, The Vincent Fairfax Family Foundation, the Hunter Medical Research Institute and the Hunter Area Pathology Service all contributed towards the costs of establishing the Hunter Community Study. The WHI program is funded by the National Heart, Lung, and Blood Institute, the US National Institutes of Health and the US Department of Health and Human Services (HHSN268201100046C, HHSN268201100001C, HHSN268201100002C, HHSN268201100003C, HHSN268201100004C and HHSN271201100004C). This work was also funded by NCI U19 CA148065-01. This research has been conducted using the UK Biobank Resource under applications 5122 and 9797.

ANECS recruitment was supported by project grants from the NHMRC [ID#339435], The Cancer Council Queensland [ID#4196615] and Cancer Council Tasmania [ID#403031 and ID#457636]. SEARCH recruitment was funded by a programme grant from Cancer Research UK [C490/A10124]. The Bavarian Endometrial Cancer Study (BECS) was partly funded by the ELAN fund of the University of Erlangen. The Hannover-Jena Endometrial Cancer Study was partly supported by the Rudolf Bartling Foundation. The Leuven Endometrium Study (LES) was supported by the Verelst Foundation for endometrial cancer. The Mayo Endometrial Cancer Study (MECS) and Mayo controls (MAY) were supported by grants from the National Cancer Institute of United States Public Health Service [R01 CA122443, P30 CA15083, P50 CA136393, and GAME-ON the NCI Cancer Post-GWAS Initiative U19 CA148112], the Fred C and Katherine B Andersen Foundation, the Mayo Foundation, and the Ovarian Cancer Research Fund with support of the Smith family, in memory of Kathryn Sladek Smith. MoMaTEC received financial support from a Helse Vest Grant, the University of Bergen, Melzer Foundation, The Norwegian Cancer Society (Harald Andersens legat), The Research Council of Norway and Haukeland University Hospital. The Newcastle Endometrial Cancer Study (NECS) acknowledges contributions from the University of Newcastle, The NBN Children’s Cancer Research Group, Ms Jennie Thomas and the Hunter Medical Research Institute. RENDOCAS was supported through the regional agreement on medical training and clinical research (ALF) between Stockholm County Council and Karolinska Institutet [numbers: 20110222, 20110483, 20110141 and DF 07015], The Swedish Labor Market Insurance [number 100069] and The Swedish Cancer Society [number 11 0439]. The Cancer Hormone Replacement Epidemiology in Sweden Study (CAHRES, formerly called The Singapore and Swedish Breast/Endometrial Cancer Study; SASBAC) was supported by funding from the Agency for Science, Technology and Research of Singapore (A*STAR), the US National Institutes of Health and the Susan G. Komen Breast Cancer Foundation.

The Nurses’ Health Study (NHS) is supported by the NCI, NIH Grants Number UM1 CA186107, P01 CA087969, R01 CA49449, 1R01 CA134958, and 2R01 CA082838. The authors thank the participants and staff of the Nurses’ Health Study for their valuable contributions as well as the following state cancer registries for their help: AL, AZ, AR, CA, CO, CT, DE, FL, GA, ID, IL, IN, IA, KY, LA, ME, MD, MA, MI, NE, NH, NJ, NY, NC, ND, OH, OK, OR, PA, RI, SC, TN, TX, VA, WA, WY. The authors assume full responsibility for analyses and interpretation of these data. The authors also thank Channing Division of Network Medicine, Department of Medicine, Brigham and Women’s Hospital and Harvard Medical School. Finally, the authors also acknowledge Pati Soule and Hardeep Ranu for their laboratory assistance. The Connecticut Endometrial Cancer Study was supported by NCI, NIH Grant Number RO1CA98346. The Fred Hutchinson Cancer Research Center (FHCRC) is supported by NCI, NIH Grant Number NIH RO1 CA105212, RO1 CA 87538, RO1 CA75977, RO3 CA80636, NO1 HD23166, R35 CA39779, KO5 CA92002 and funds from the Fred Hutchinson Cancer Research Center. The Multiethnic Cohort Study (MEC) is supported by the NCI, NHI Grants Number CA54281, CA128008 and 2R01 CA082838. The California Teachers Study (CTS) is supported by NCI, NIH Grant Number 2R01 CA082838, R01 CA91019 and R01 CA77398, and contract 97-10500 from the California Breast Cancer Research Fund. The Polish Endometrial Cancer Study (PECS) is supported by the Intramural Research Program of the NCI. The Prostate, Lung, Colorectal, and Ovarian Cancer Screening Trial (PLCO) is supported by the Extramural and the Intramural Research Programs of the NCI.

Melbourne Collaborative Cohort Study (MCCS) cohort recruitment was funded by VicHealth and Cancer Council Victoria. The MCCS was further augmented by Australian National Health and Medical Research Council grants 209057, 396414 and 1074383 and by infrastructure provided by Cancer Council Victoria. Cases and their vital status were ascertained through the Victorian Cancer Registry and the Australian Institute of Health and Welfare, including the National Death Index and the Australian Cancer Database.
